# Metagenome-assembled genomes from *Monte Cristo* Cave (Diamantina, Brazil) reveal prokaryotic lineages as functional models for life on Mars

**DOI:** 10.1101/2020.07.02.185041

**Authors:** Amanda G. Bendia, Flavia Callefo, Maicon N. Araújo, Evelyn Sanchez, Verônica C. Teixeira, Alessandra Vasconcelos, Gislaine Battilani, Vivian H. Pellizari, Fabio Rodrigues, Douglas Galante

## Abstract

Although several studies have explored microbial communities in different terrestrial subsurface ecosystems, little is known about the diversity of their metabolic processes and survival strategies. The advance of bioinformatic tools is allowing the description of novel and not-yet cultivated microbial lineages in different ecosystems, due to the genome reconstruction approach from metagenomic data. The recovery of genomes has the potential of revealing novel lifestyles, metabolic processes and ecological roles of microorganisms, mainly in ecosystems that are largely unknown, and in which cultivation could be not viable. In this study, through shotgun metagenomic data, it was possible to reconstruct several genomes of cultivated and not-yet cultivated prokaryotic lineages from a quartzite cave, located in Minas Gerais state, Brazil, which showed to possess a high diversity of genes involved with different biogeochemical cycles, including reductive and oxidative pathways related to carbon, sulfur, nitrogen and iron. Tree genomes were selected, assigned as *Truepera sp*., *Ca. Methylomirabilis sp*. and *Ca. Koribacter sp*. based on their lifestyles (radiation resistance, anaerobic methane oxidation and potential iron oxidation) for pangenomic analysis, which exhibited genes involved with different DNA repair strategies, starvation and stress response. Since these groups have few reference genomes deposited in databases, our study adds important genomic information about these lineages. The combination of techniques applied in this study allowed us to unveil the potential relationships between microbial genomes and their ecological processes with the cave mineralogy, as well as to discuss their implications for the search for extant lifeforms outside our planet, in silica- and iron-rich environments, especially on Mars.

## Introduction

Caves are an example of the terrestrial dark settings that harbor an extensive microbial biomass, biodiversity and geochemical processes, which generates noteworthy morphological and molecular biosignatures, such as the those produced by the recognition of metabolic potentials though specific gene detection (Boston et al., 2001; Hershey and Barton, 2018; Tetu et al., 2013; Wiseschart et al., 2019). Few studies describing the microbial diversity of cave ecosystems have detected the metabolic potential of these microorganisms. In fact, metabolic potential-based recognition of cave microbiome is one of the least scientifically explored strategies (Riquelme et al., 2017). These few studies pointed that, in the absence of light, the microbial community relies on diverse chemolithotrophic metabolisms, as iron oxidation, sulfide oxidation, sulfate reduction, hydrogen oxidation, methanotrophy and methanogenesis (e.g. Osburn et al., 2014; Riquelme et al., 2017; Simkus et al., 2016; Wiseschart et al., 2019). Microbial metabolism reported in cave ecosystems from Thailand and Sweden achieved in the Manao-Pee cave and the Tjuv-Ante’s cave, respectively, shed light on the value of metabolic potentials for the understanding of microbial diversity, metabolic versatility and adaptation to the extreme conditions of terrestrial subsurface environments (Mendoza et al., 2016; Wiseschart et al., 2019).

Also, another approach aiming the understanding of microbial dynamics from cave environments is the identification of direct relationship between microorganisms and cave mineralogy, and the corresponding physicochemical signature originated from this interaction. Sauro et al. (2018) suggested that some putative microbial metabolic activities play a role in dissolution and reprecipitation of silica under cave conditions. Additionally, many studies addressed the role played by iron-oxidizing bacteria for the precipitation within Fe-biomineralized filaments and stalks in iron-rich caves, making these bioprecipitate minerals a significant inorganic biosignature (Baskar et al., 2008; Chan et al., 2016; Florea et al., 2011).

Two innovative applications of the metabolic potential and diversity in caves ecosystems might be in astrobiology, contributing to elucidate the potential metabolisms in extraterrestrial analogous oligotrophic systems, such as subsurface environments on Mars (Hays et al., 2017; Ortiz et al., 2014). Because of the hostile surface conditions on Mars (e.g. the incidence of intense UV-C radiation) the subsurface environments may offer one of the only accesses to recognizable biosignatures of extant lifeforms (Boston et al., 2001). Also, caves on Earth are particularly interesting analogues because of the better preservation potential of biosignatures over long periods (< 10^4^ years for DNA and <10^6^ years for proteins; Bada et al., 1999), once they may stand under the same physicochemical conditions for long periods (up to millennia time scale; Hershey and Barton, 2018).

Another possible application of specific metabolic potentials for astrobiological purposes is related to the characteristics of Mars’ surface. The Martian environment, either superficial or on the subsurface, presents many stressors and challenges for life. The diversity of metabolisms found on caves, especially those that allow the exploration of non-conventional sources of energy and redox potential under extreme conditions, make them promising analogue environments for Mars, and should be further explored. For example, metabolic potentials based on methane and iron may be of great interest, once these elements have been identified on Mars’ surface (e.g. Johnson et al., 2016; Rieder et al., 1997; Singer et al., 1979). Recently, sub-annually changes in atmospheric concentration of methane were detected by the Curiosity rover in the area of Gale Crater (Hu et al., 2016) and life was not ruled out among the hypotheses proposed to explain the origin of Martian methane. In addition, these methane-producing regions could harbor a methanotrophic biota.

Thus, understanding the ecological spectra and adaptation aspects of the metabolic diversity found on caves under oligotrophic settings may be important for i) understanding the several chemolithotrophic metabolisms, including those that better flourish in areas with low carbon offer; ii) expand our knowledge concerning potential areas for habitability; and, again, iii) be useful for life prospection in our neighboring planets and other surfaces.

The Brazilian territory harbors several karst ecosystems that are still poorly explored regarding their microbial diversity. The *Monte Cristo* cave consists of a quartzite cave, presenting aphotic and oligotrophic conditions. It is located in Serra do Espinhaço mountain ridge, near the city of Diamantina, in the State of Minas Gerais, Brazil, an iron-rich area formed by Proterozoic metasedimentary rocks (Vasconcelos, 2014).

Based on the exposed before, the main goal of the present research was, through the genome reconstruction of cultivated and not-yet cultivated prokaryotic lineages, to reveal the metabolic diversity and molecular survival strategies under the geological environment of the *Monte Cristo* cave, being one of the first studies that recovered metagenome-assembled genomes from a karst ecosystem. By combining genome reconstruction from shotgun metagenomic data, synchrotron-based X-ray diffraction and physicochemical characterization, this study has implications for the better understanding of microbial functions in quartzite karstic ecosystems and assess the strategies of life to thrive in extreme conditions. Also, it may be useful for suggestions of potential metabolic processes to be explored on Mars’ subsurface, serving as a basis for astrobiological studies.

## Methodology

### Study area and geological setting

The *Monte Cristo* Cave is located in the eastern border of Meridional Serra do Espinhaço, at the locality known as Extração, in southeast Diamantina, state of Minas Gerais, Brazil (Figure 1). The Serra do Espinhaço comprises a north-south oriented mountain range, extending from the state of Minas Gerais to the state of Bahia, totalizing an extension of 1000 km in Southeast Brazil. Due to its extension and topographic variation, the climate and soil type are varied as well.

**Figure 1.**
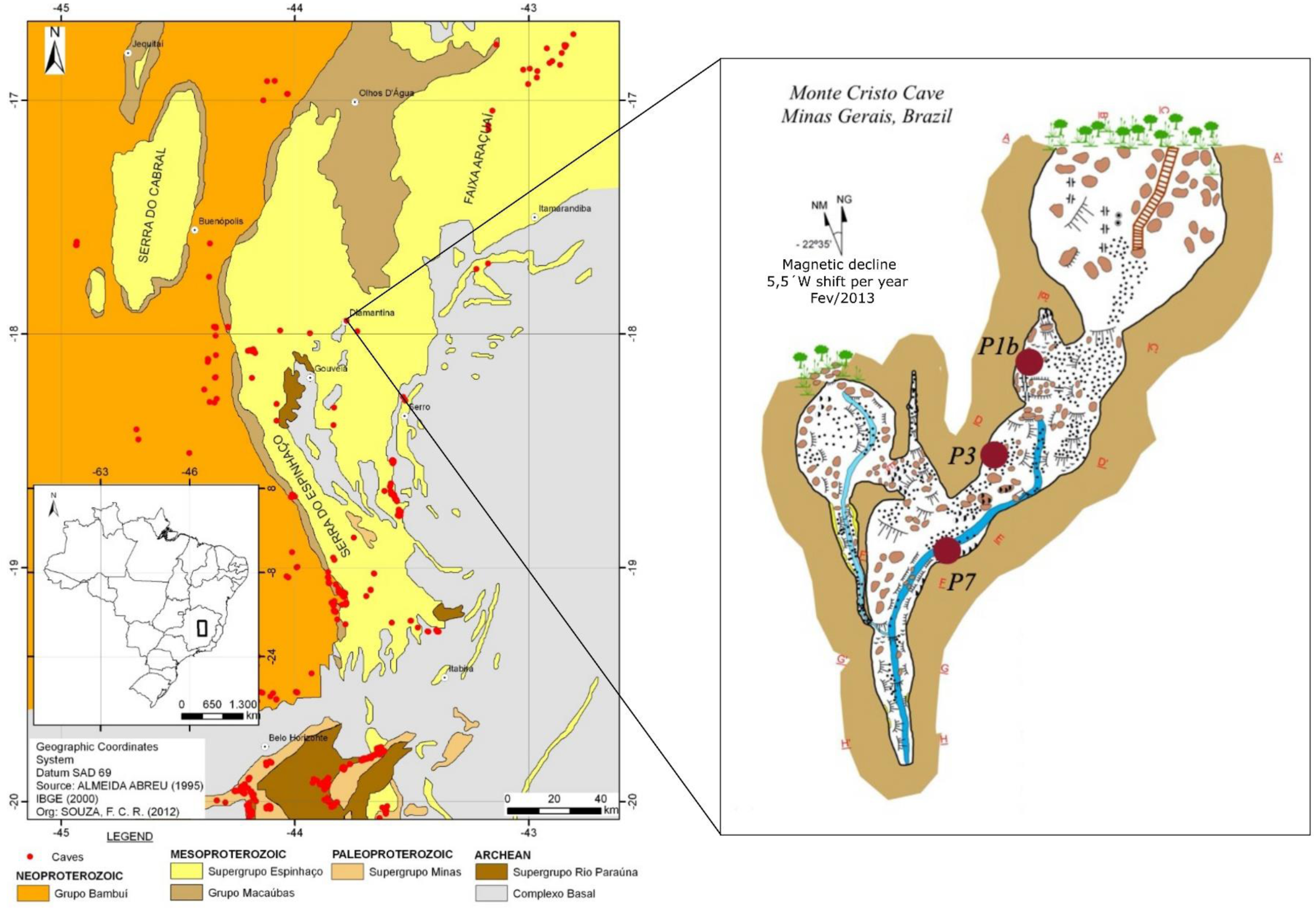
Geological context of Meridional Serra do Espinhaço mountain range. *Monte Cristo* Cave, located near the city of Diamantina, State of Minas Gerais. Monta Cristo cave map was adapted from Souza (2014).

The geological context of the studied area corresponds to the Espinhaço Supergroup (Figure 1). This unit comprises sedimentary to metasedimentary (greenschist facies) rocks deposited in an intracratonic rift during the Paleo-to Mesoproterozoic. At the basal portion, the supergroup includes the diamondiferous metaconglomerates and continental fluvial, fan-deltaic, and lacustrine quartzites represented by the São João da Chapada and Sopa-Brumadinho formations (Alkmim et al., 2006; Dussin and Dussin, 1995), both representing the Guinda Group (Knauer and Schrank, 1993). Upwards, the Espinhaço Supergroup includes the aeolian quartzites of Galho do Miguel Formation, overlaid by the Conselheiro Mata Group, which includes five formations, all deposited under marine conditions, and represented by metasiltstones, metadolomite and quartzite deposits (Almeida Abreu and Pflug, 1994; Silva and Chaves, 2012).

The *Monte Cristo* Cave was developed in the metaconglomerates of Sopa-Brumadinho Formation. This unit includes, besides the diamantiferous metaconglomerades, alkaline to acid metavolcanic rocks deposited in three, 20 to 50 meters-thick cycles (Abreu, 1995; Chula et al., 2013), associated to a peak in extensional tectonic activity during Late Paleoproterozoic. Schöll and Fogaça (1979) proposed the subdivision of Sopa-Brumadinho Formation in three members, and (Almeida Abreu and Pflug, 1994) named them, being from the base upwards, the phyllites and quartz-phyllites of Datas Member; the Fe-rich quartzites and pure quartzites of Caldeirões Members; and the phyllites, metasiltstones, quartzites, and metabreccias of Campo Sampaio Member.

At the area of *Monte Cristo* Cave, the lithostratigraphic context is interpreted as the Caldeirões Members. Locally, 15 meters-thick level of pelites occurs in angular unconformity with medium to coarse psammite levels intercalated with conglomerates deposits of the São João da Chapada Formation. Those conglomerates can be; i) clast-supported, presenting greenish pelitic matrix and pebble to boulder clasts of quartzites, metavolcanic rocks, and banded iron formations nature; and ii) sand-supported, presenting pebbles and boulders of quartzite and, eventually, banded iron formation (Chaves, 1997; Silva and Chaves, 2012). Quartzites outcrops associated with monolithic conglomerate lens, sericite-matrix orthoconglomerates, quartz-matrix paraconglomerates, as well dikes and sills of alkaline and metalkaline rocks were also identified in the cave area.

Structural geology analysis pointed for the occurrence of structures with plane-axial character, represented by schistosity and cleavage in areas with little deformation, while in shear zones mylonitic foliation may occur associated with anastomosed features (Rosière et al., 1994). In high deformed areas, ramps and flats associated with thrust faults, indicating a brittle to ductile-brittle regime. Litostructural analysis also pointed for structural lineaments as the important factor, although not the only one, for caves developing in the Extração area, including the *Monte Cristo* Cave (Souza and Salgado, 2014).

### Sampling procedure

Samples were collected in May 2018 in three localities of the *Monte Cristo* cave (18° 17.822’S, 43° 33.511’W), Minas Gerais state, Brazil. The sampling was performed aseptically on the surface of the cave wall (first ∼5 cm) and included three sample types: dry rock (P1B), dry saprolite (P3) and wet saprolite (P7) (Table 1). Samples were stored for molecular analysis in sterile packs and placed at -20°C. The sample collected at point P1B (Figure 2A) consists of fragments of the purplish rock from the cave wall. At point P3 (Figure 2B), the cave wall was also sampled and consisted of a yellowish-colored saprolite, in which small hard nodules of millimeter size were formed. At point P7 (Figure 2C), the sample comprised a brownish saprolite above the interface between sediments and a small stream, being the only wet sample collected.

**Table 1.** Description physicochemical parameters and mineralogy of samples from *Monte Cristo* cave, the prevalent microbial genomes, and the number of reads obtained from shotgun metagenomics. The elemental concentrations are expressed in values of ppm (mg/kg or mg/L) related to each 1g of solubilized sample.

| Point | Description | pH | Conductivity ($\mu S/cm$) | Element concentration | | | | | | Mineralogy (*) | Total reads | Filtered reads | Prevalent microbial genomes |
| --- | --- | --- | --- | --- | --- | --- | --- | --- | --- | --- | --- | --- | --- |
|  |  |  |  | Cu (ppm) | Fe (ppm) | Mn (ppm) | Ni (ppm) | S (ppm) | Zn (ppm) |  |  |  |  |
| 1B | dry sample, consolidated substrate<br>rock | 5.59 | 1424 | 0.133 | 114.58 | 22.683 | <QL | 112.25 | 1.83 | quartz (SiO<sub>2</sub>),<br>variscite (AlPO<sub>4</sub>(H<sub>2</sub>O)<sub>2</sub>),<br>rutile (TiO<sub>2</sub>),<br>phlogopite (KMg<sub>3</sub>(Si<sub>3</sub>Al)O<sub>10</sub>F<sub>2</sub>)<br>hematite (Fe<sub>2</sub>O<sub>3</sub>) | 53,774,058 | 50,429,810 | Actinobacteriota,<br>Proteobacteria,<br>Firmicutes,<br>Bacteroidota,<br>Patescibacteria,<br>Deinococcota |
| 3 | dry sample, unconsolidated substrate<br>saprolite | 4.41 | 194 | <QL | 176.43 | 4.119 | 0.104 | 54.087 | 0.440 | jarosite (K<sub>2</sub>Fe<sub>6</sub>(OH)<sub>12</sub>(SO<sub>4</sub>)<sub>4</sub>),<br>rutile (TiO<sub>2</sub>),<br>phlogopite (KMg<sub>3</sub>(Si<sub>3</sub>Al)O<sub>10</sub>F<sub>2</sub>) and<br>quartz (SiO<sub>2</sub>). | 3,453,312 | 3,232,448 | *Mycobacterium sp.* |
| 7 | wet sample, unconsolidated substrate (near a small stream)<br>saprolite | 7.15 | 28 | 0.014 | 227.99<br>3 | 6.494 | <QL | 1.012 | 0.0297 | quartz (SiO<sub>2</sub>),<br>muscovite (KAl<sub>2</sub>(Si<sub>3</sub>Al)O<sub>10</sub>(OH, F)<sub>2</sub>), kaolinite (Al<sub>2</sub>Si<sub>2</sub>O<sub>5</sub>(OH)<sub>4</sub>),<br>rutile (TiO<sub>2</sub>). | 55,237,758 | 51,938,168 | Nitrososphaeria,<br>Dadabacteria,<br>Chloroflexota,<br>Acidobacteriota,<br>Nitrospirota,<br>Dormibacterota,<br>Methylomirabilota |

Quantification limit (QL) for the elements Cu and Ni is approximately 0.01 ppm for sample.
1% = 10,000 ppm (mg/kg or mg/L).
(\*) based on X-ray diffraction analysis.

**Figure 2.**
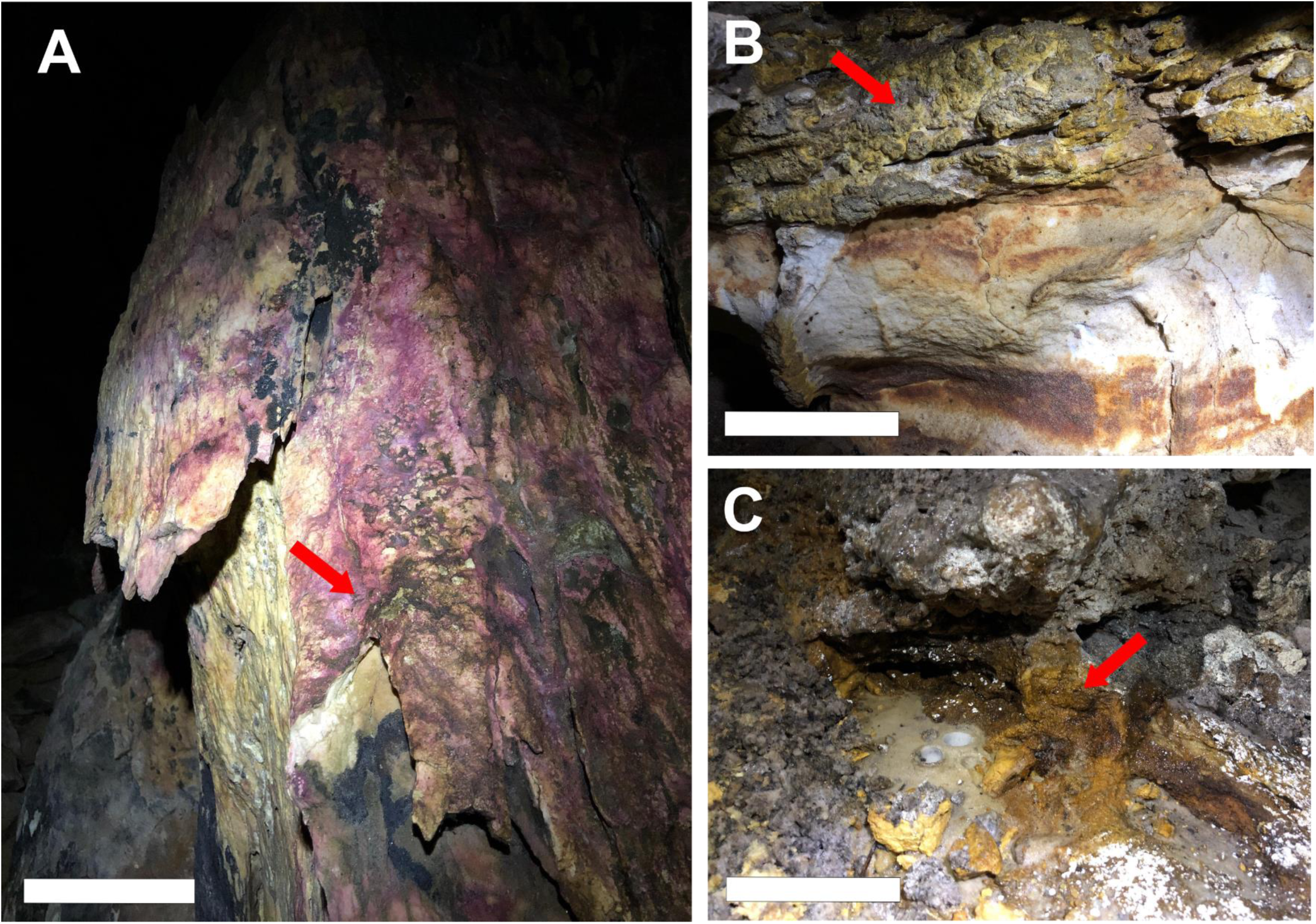
Sampled points at Monte Cristo Cave. The red arrows point to the exact point of extraction of each sample. A) Point P1B, showing the hard purplish crust in the cave wall. Scale bar = 30 cm; B) Point P3, consisting of yellowish-colored crust with small nodules. Scale bar = 10 cm; C) Point P7, brownish crust in the interface between water and sediments.

### Physicochemical and mineralogical analysis

The wet substrate sample was previously dried and all samples were powdered in an agate mill. For pH and conductivity measurements, the samples were mixed with deionized water. The quantitative elemental characterization was performed at Chemistry Institute of University of São Paulo using the inductively coupled plasma-atomic emission spectrometry (ICP-OES) to measure aliquots previously opened with 10% HNO3. The characterization of mineralogical content was performed by X-ray diffraction at Brazilian Synchrotron Light Laboratory and its User Chemistry Laboratory. The measurements were performed at XRD1 beamline, using the geometry of Debye-Scherrer, excitation energy of 12 keV and the Mythen detector in a full range of 2 theta (5 to 120 degrees). The diffraction data was measured at 12 keV and the crystalline phases were indexed according to The International Center for Diffraction Data (ICDD).

### DNA extraction and metagenomic sequencing

DNA was extracted using Power Soil DNA extraction kit (Qiagen) following the instructions provided by the manufacturer. The quantification was performed with Qubit and Qubit® dsDNA HS Assay Kit (Life Technologies, USA). Total DNA samples were submitted to shotgun metagenomic sequencing using Illumina Hiseq 2×150 pb platform. Sequencing was supported by The Deep Carbon Observatory’s Census of Deep Life initiative and was performed at the MBL (Woods Hole, MA, USA).

### Metagenomic assembly and genomic reconstruction

Metagenomic data was processed using anvi’o v. 5 pipeline, following the workflow described by Eren et al. (2015). First, reads were quality filtered through -iu-filter-quality-minoche, and then co-assembled using MEGAHIT v. 1.0.2. (Li et al., 2015), discarding contigs smaller than 1000 bp. Reads from each sample were mapped to the co-assembly using bowtie2 with default parameters (Langmead and Salzberg, 2012), and then it was generated a contig database through -anvi-gen-contigs-database. Prodigal (Hyatt et al., 2010) was applied to predict the open reading frames (ORFs). HMMER v. 3.1b2 (Finn et al., 2011) was utilized to identify single-copy bacterial and archaeal genes, and -anvi-run-ncbi-cogs was applied to gene annotation using Clusters of Orthologous Groups (COGs) database (December 2014 release with blastp v2.3.0+ (Altschul et al., 1990). GhostKOALA (genus_prokaryotes) (Kanehisa et al., 2016) was employed for functionally and taxonomically annotations of the predicted protein sequences. The -anvi-profile was used to profile individual BAM files, with a minimum contig length of 4 kbp. Genome binning was performed using CONCOCT (Alneberg et al., 2013) through ‘anvi-merge’ program with default parameters. Manually refined bins using ‘anvi-refine’ were adopted, and completeness and contamination were estimated using ‘anvi-summarize’. Bins were quality checked through CheckM v. 1.0.7 (Parks et al., 2015), which is based on the representation of lineage-specific marker gene sets. Bins were taxonomically classified based on genome phylogeny using GTDB-Tk (Chaumeil et al., 2020). Raw reads are deposited in NCBI’s Genbank under the Bioproject PRJNA642310, and the Biosamples IDs SAMN15393797, SAMN15393798, SAMN15393799.

### Taxonomic and functional annotation of metagenome-assembled genomes (MAGs)

Bins were classified as high-quality draft (>90% complete, <5% contamination), medium-quality draft (>50% complete, <10% contamination) or low-quality draft (<50% complete, <10% contamination) metagenome assembled-genome (MAG), accordingly with genome quality standards suggested by (Bowers et al., 2017). Annotation of MAGs was performed using prokka (Seemann, 2014). Proteins were compared to sequences in the KEGG Database through GhostKOALA (genus_prokaryotes) (Kanehisa et al., 2016) and in SEED Subsystem through RASTtk (Brettin et al., 2015) in order to reveal metabolic potential of MAGs. FeGenie tool (Garber et al., 2020) was employed to annotate genes related to iron metabolism in MAGs and MetabolismHMM tool (https://github.com/elizabethmcd/metabolisHMM) to annotate genes related to the sulfur, nitrogen and carbon metabolisms.

Pangenome analysis was performed using the anvi’o v. 5 pipeline to conduct genome comparison and gene clusters identification of three selected MAGs: DC_MAG_00005, DC_MAG_00016 and DC_MAG_00021. These MAGs were selected according to their quality, metabolic and survival lifestyles: resistance to ionizing radiation (DC_MAG_00005), anaerobic methane oxidation (DC_MAG_00021) and potential iron oxidation (DC_MAG_00016). Further, there are few genomes of these taxonomic groups in databases and novel genomes might provide important information regarding the molecular strategies of these lineages. The pangenome analysis of DC_MAG_00005 (*Truepera sp*.) was performed by the comparison with the genome of *Truepera radiovictrix* (DSM17093) and the MAG assigned as *Truepera sp*. (GCA_002239005.1_ASM223900), the only genomes of the family Trueperaceae deposited in NCBI database. Also, the DC_MAG_00016 (*Ca. Koribacter*) was compared with the *Ca. Koribacter versatilis* (NC_008009.1), the only genome of this genus deposited in NCBI database. For DC_MAG_00021 (*Ca. Methylomirabilis*), *Ca. Methylomirabilis oxyfera* (FP565575.1) and *Ca. Methylomirabilis limnetica* (NZ_NVQC01000028.1) were selected. The Average Nucleotide Identity calculator (ANI) and Average Amino acid Identity calculator (AAI) available in Kostas Lab website (http://enve-omics.ce.gatech.edu/) were applied to compare our MAGs with the reference genomes.

## Results

### Physicochemical and mineralogical analysis

Average values from triplicates of physicochemical characterization (pH values and conductivity), elemental concentration (in ppm) and the mineralogical characterization are summarized in Table 1. P7 presented the highest value (227.993 ppm) in comparison with the other sampled points. P1B and P3 presented the highest values for S (112.25 and 54.087 ppm, respectively) in comparison with P7 (1.012 ppm). In addition, P1B was also rich in Mn (22.683 ppm). The other measured elements (Zn, Cu and Ni) presented low values, being below the detection limit of the equipment at some points.

The main observed phase in the diffractogram of all samples is Quartz (SiO2, ICDD 00-046-1045). 2theta is presented from 5 – 80 ° and the insets present secondary phases for each material. Secondary reflections (Figure 3) were identified in P1B as Variscite (AlPO_4_ (H_2_ O)_2_, ICDD-01-070-0310), Rutile (TiO_2_, ICDD 00-034-0180), Phlogopite (KMg_3_ (Si_3_ Al)O_10_ F_2_, ICDD 01-076-0629) and Hematite (Fe_2_O_3_, ICDD-01-073-0603). For P3, samples presented Jarosite (K(Fe_3_ (SO_4_)_2_ (OH)_6_, ICDD 01-076-0629), Rutile (TiO_2_, ICDD 00-034-0180) and Phlogopite (KMg_3_ (Si_3_Al)O_10_F_2_, ICDD 01-076-0629). For P7, it was possible to identify Muscovite (KAl_2_ (Si_3_Al)O_10_ (OH, F)_2_, ICDD-00-006-0263), Kaolinite (Al_2_Si_2_O_5_ (OH)_4_, ICDD-01-075-0938) and Rutile (TiO_2_, ICDD 00-034-0180). The relative intensity of reflections is associated with the proportions and/or crystallinity degree of each phase on the mixture. Some phases are represented by a single, most intense, indexed-peak, in a consequence from the way the structure reflects, so it means the analysis just suggests the presence of those minerals (rutile, hematite, kaolinite and phlogopite). However, minerals such as jarosite, variscite and muscovite are represented at least by three peaks, which suggest these materials are really part of the sample. It is important to mention there are still some extra peaks relative to less intense phases not identified up to now.

**Figure 3.**
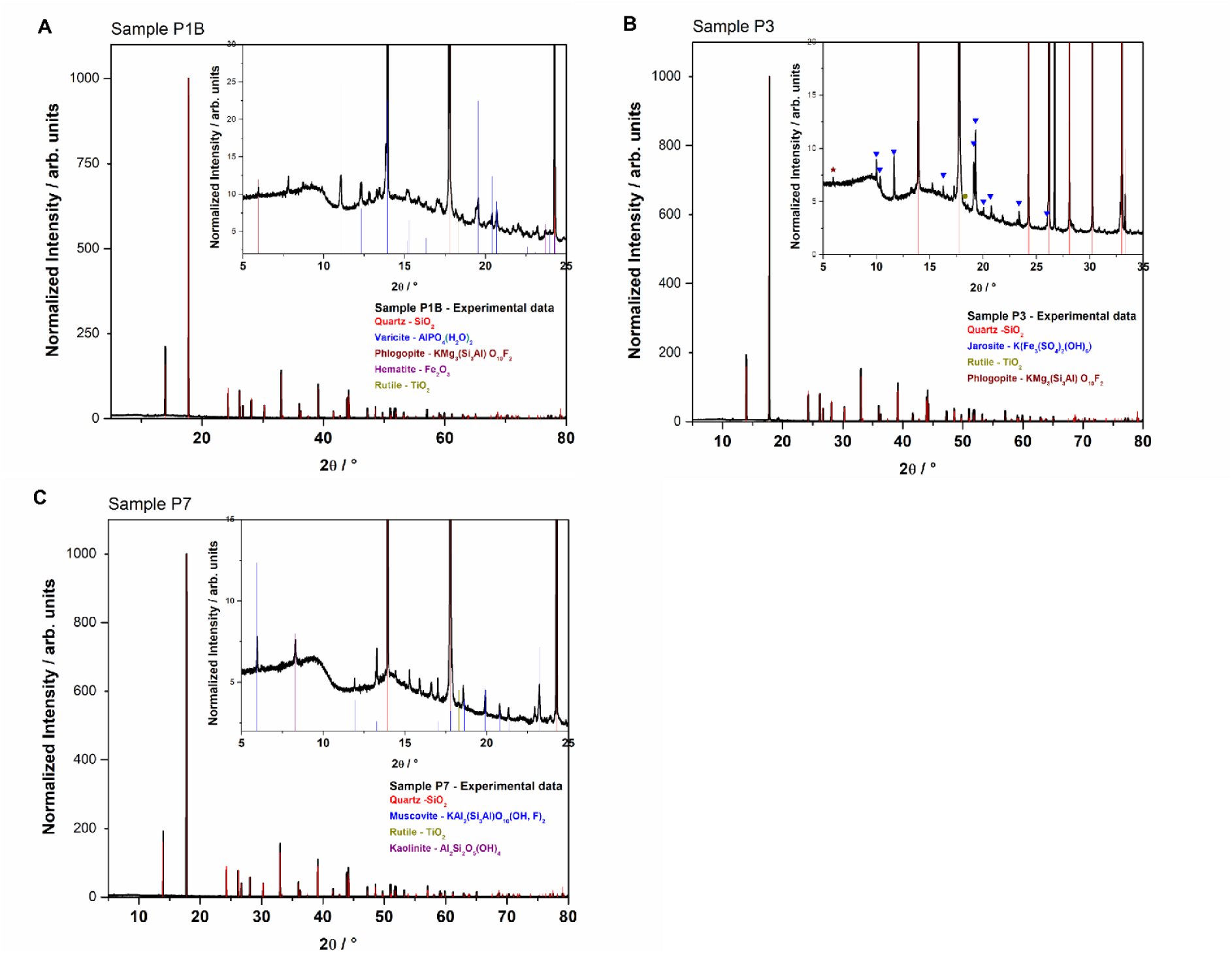
Diffractograms of the samples collected in A) P1B, showing the mineral phases quartz, varicite, phlogopite, hematite and rutile; B) P3, with quartz, jarosite, rutile and phlogopite, and 3) P7, presenting quartz, muscovite, rutile and kaolinite.

### Quality filtering and assembly

A total of 112,465,128 raw paired-end reads were obtained, distributed among three cave samples, in which 105,600,426 reads were filtered by quality and then used for co-assembly (Table 1). Co-assembly resulted in 302,685 contigs with >1000 pb and a N50 of 4,212 bp, with the longest contig containing 863,531 bp. 1,135,436 genes were detected in the contigs through prodigal, and 10,439 HMM hits for Archaea and 15,170 for Bacteria.

### Taxonomic classification of metagenome-assembled genomes

Using anvi’o, a total of 125 MAGs were recovered from *Monte Cristo* cave, in which 13 were defined as high-quality drafts (>90% completeness, <5% contamination) and 48 as medium-quality drafts (>70% completeness, <5% contamination). The general view of MAGs is represented in Figure 4A and Supplementary Table 1, including the values of completeness, contamination, GC content, total reads mapped and number of SNVs. The 61 high-medium-quality genome drafts were selected for taxonomical and functional annotation. Among these, the majority was assigned within Actinobacteria (21.3%) and Proteobacteria (19.7%), and the others as Acidobacteriota (8.2%), Crenarchaeota (6.5%), Bacteroidota (6.5%), Firmicutes (6.5%), Nitrospirota (6.5%), Chloroflexota (4.9%), Patescibacteria (4.9%), Bipolaricaulota (3.3%), Dadabacteria (1.6%), Deinococcota (1.6%), Methylomirabilota (1.6%) and 6.5% as unclassified bacterial phylum (Figure 4C). In general, the majority of MAGs were obtained from sample P1B and P7. Only 1 MAG was recovered from sample P3, classified as *Mycobacterium*. The MAGs from sample P1B were mainly assigned within Actinobacteriota, Proteobacteria, Firmicutes, Bacteroidota, Patescibacteria and Deinococcota, whereas those from P7 were mainly classified as Nitrososphaeria, Chloroflexota, Acidobacteriota, Nitrospirota, Dormibacterota and Methylomirabilota (Figure 4B).

**Figure 4.**
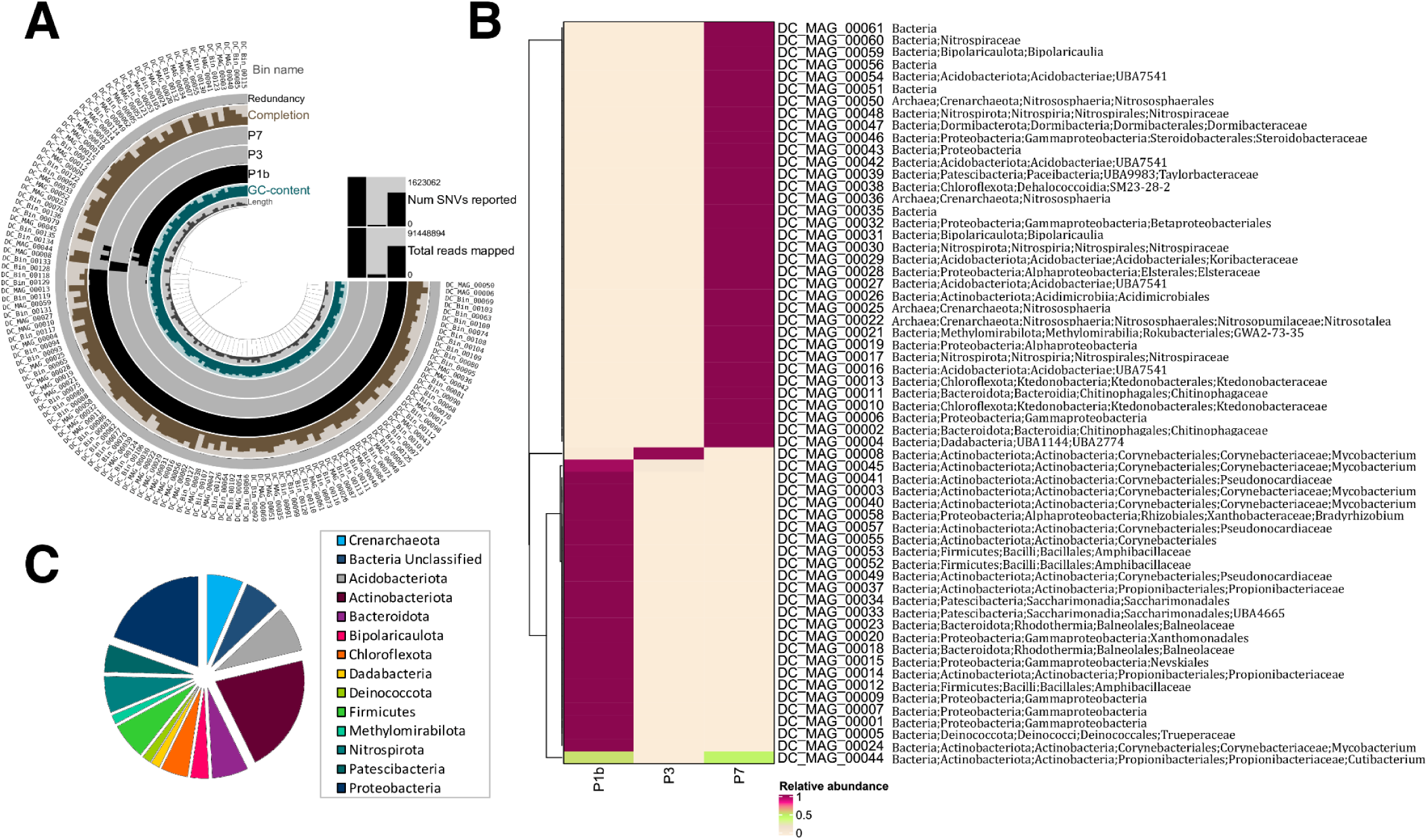
General results of the metagenome-assembled genomes are represented in A, including the mean coverage for each bin, their completeness and redundancy, the total reads mapped and number of SNVs reported. The taxonomy classification of the 61 medium- and high-quality genomes, and the relative abundances of the contigs for each sample are represented in B. The taxonomy is represented until the deepest classified level according to GTDB-Tk. The relative proportions of each phyla related to the 61 MAGs are visualized in C.

### Metabolic potential of metagenome-assembled genomes

Functional annotations of the 61 high-medium-quality MAGs were performed in order to investigate the metabolic potential related to carbon, nitrogen, sulfur and iron cycles, and linked to hydrogenotrophic microorganisms (Supplementary Figure 1) (Figure 5). Iron oxidation genes were detected among 10 MAGs, including members from Nitrospiraceae, Bipolaricaulia, Steroidobacteriaceae, Koribacteraceae, unclassified Proteobacteria and Acidobacteriae. Hydrogenases were found in 17 MAGs, including members from Nitrospiraceae, Bipolaricaulia, Amphibacillaceae, Pseudocardiaceae, *Mycobacterium*, Taylorbacteraceae, Balneolaceae, Xanthomonadales, Nevskiales, Propionibacteraceae, Ktedonobacteraceae and unclassified Gammaproteobacteria.

**Figure 5.**
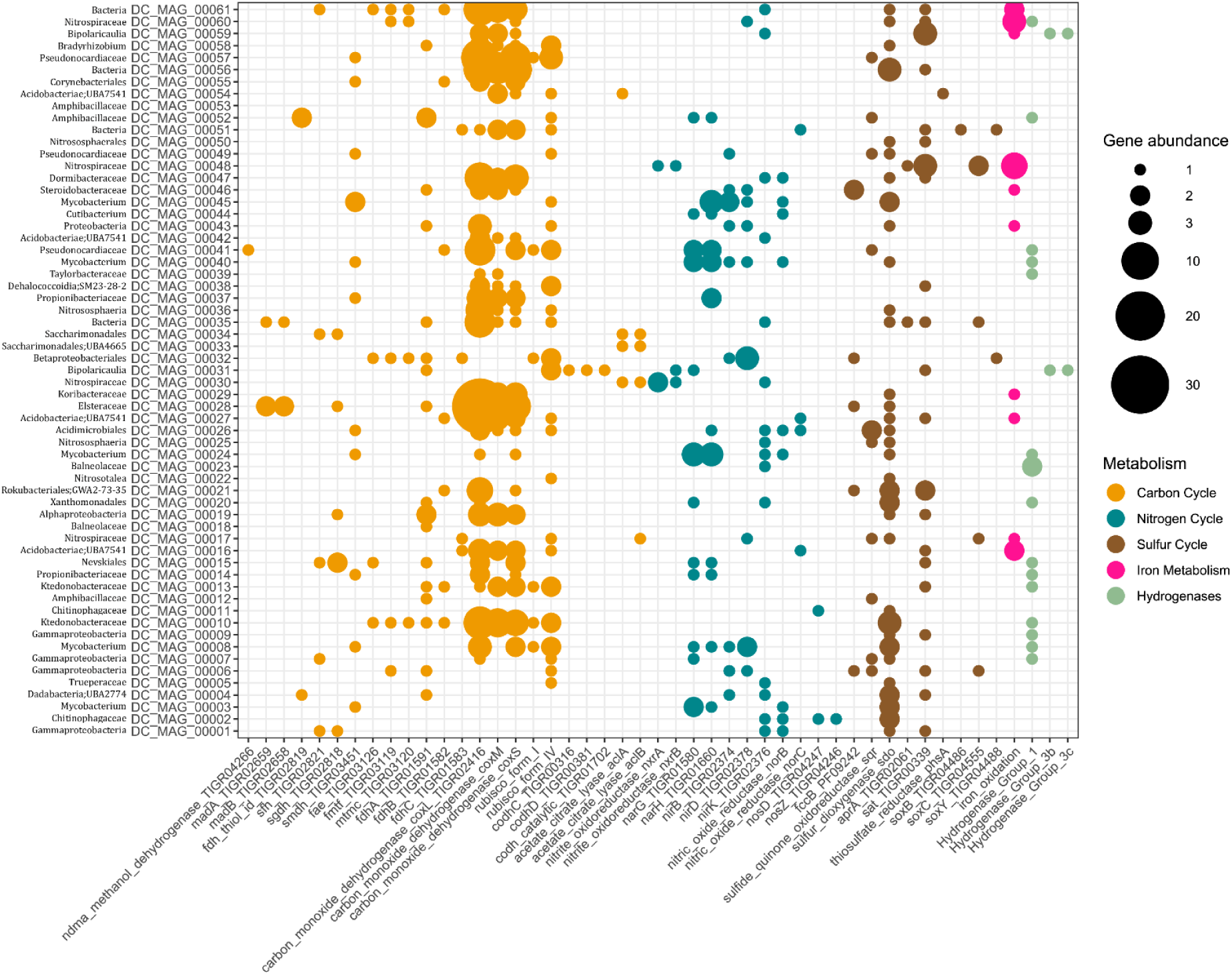
Genes related to metabolic potential of the 61 medium- and high-quality genomes are represented as absolute abundance. Genes involved with the metabolisms of carbon, nitrogen and sulfur, as well as hydrogenases, were selected by MetabolisHMM tool. Genes involved with the iron cycle were selected by the FeGenie tool. The taxonomy for the deepest classified level according to GTDB-Tk is represented for each MAG.

Regarding nitrogen cycle, several of our MAGs exhibited genes related to denitrification, such as dissimilatory nitrate reduction (*narG* and *narH*), nitrite reduction (*nirB, nirD* and *nirK*), nitric oxide reduction (*norB* and *norC*) and nitrous oxide reduction (*nosD* and *nosZ*) pathways. Nitrification genes (*nxrA* and *nxrB*) were only detected in three MAGs, classified as Nitrospiraceae and Bipolaricaulia. Genes related to sulfur oxidation (*sox* system) were detected among 6 MAGs, including the *soxB* gene in MAG00051 (unclassified Bacteria), *soxC* in MAG00048 (Nitrospiraceae), MAG00006 (Gammaproteobacteria), MAG00017 (Nitrospiraceae) and MAG00035 (unclassified Bacteria), and *soxY* in MAG00051 and MAG00032 (Betaproteobacteriales). Other genes related to sulfur oxidation were also detected in our MAGs, such as sulfur dyoxigenase (*sdo*) and sulfide quinone oxidoreductase (*sqo*). Sulfate reduction genes were found in 21 MAGS for *sat* and in 2 MAGs for *aprA*.

Genes involved with three CO_2_ -fixation pathways were identified among MAGs. The Calvin-Benson-Bassham (CBB) cycle (RuBisCO type I) was detected among 5 MAGs including members from *Mycobacterium*, Ktedonobacteraceae Betaproteobacteria and Pseudonocardiaceae. ATP-citrate lyase, the defining enzyme of the Arnon–Buchanan reductive tricarboxylic acid (rTCA) cycle, was found in 5 MAGs classified within Nitrospiraceae, Patescibacteria and Acidobacteria. The CO-methylating acetyl-CoA synthase, the crucial enzyme for the Wood-Ljungdahl (WL) pathway, was found in one MAG assigned to Bipolaricaulia. Our MAGs also exhibited genes related to methane metabolism, mainly methylotrophy. Three subunits of formate dehydrogenase gene (*fdhABC*) were found in several MAGs, whereas methanol dehydrogenase (*mdh*) was detected only in DCMAG00041 (Pseudocardiaceae).

### Genome comparison and survival strategies of DC_MAG_00005, DC_MAG_00016 and DC_MAG_00021

Three MAGs were selected to perform pangenomic analysis and annotation of survival strategies genes, including DNA repair, starvation and stress response processes. Reference genomes available in NCBI database were selected for comparison. DC_MAG_00005 (completeness 95%, contamination 0%), assigned as *Truepera sp*., was selected due to its known capacity of radiation resistance. DC_MAG_00016 (completeness 90%, contamination 0%), classified as *Ca. Koribacter*, was selected by its chemolithotrophic metabolism, and its genes related to iron oxidation. DC_MAG_00021 (completeness 88%, contamination 0.7%), classified as *Ca. Methylomirabilis*, it was selected due to its nitrite-dependent anaerobic methane oxidation capacity.

The pangenome analysis of DC_MAG_00005 (*Truepera sp*.) resulted in a total of 702 gene clusters that was considered as the core genome, whereas 587 were accessory (present in two genomes) and 5,509 were singletons (present in only one genome) (Figure 6). The pangenome analysis of DC_MAG_00016 (*Ca. Koribacter*) resulted in 645 gene clusters as the core genome and 7,009 gene clusters as singletons. For DC_MAG_00021 (*Ca. Methylomirabilis*), the pangenome resulted in 407 gene clusters as core genome, 1,238 as accessory and 4,244 as singletons. The results indicate an open pangenome for these groups, since many singletons and/or accessory genes are present. In addition, our MAGs exhibited <70% of ANI and AAI in comparison with their reference genomes, which suggests they belong to a distinct species, or even to a distinct family.

**Figure 6.**
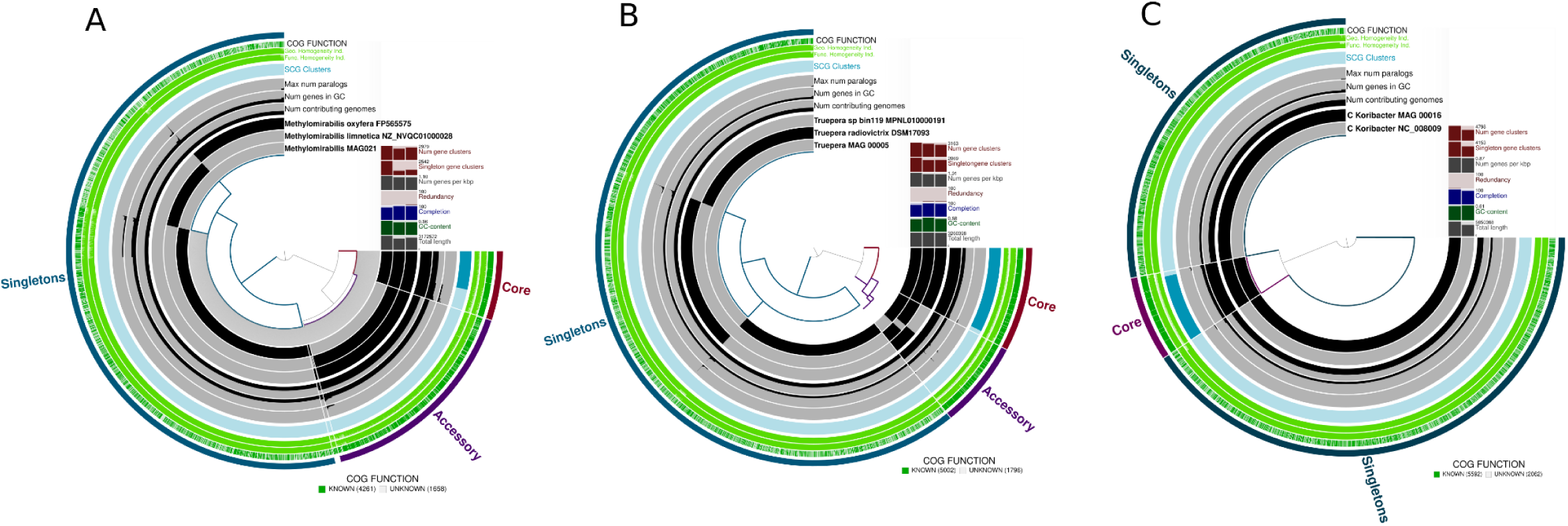
Pangenomic comparison of DC_MAG_00021 (*Ca. Methylomirabilis sp*.) (A), DC_MAG_00005 (*Truepera sp*.) (B) and DC_MAG_00016 (*Ca. Koribacter sp*.) (C), with their reference genomes deposited in NCBI. General information about the number of gene clusters, singleton gene clusters, number of genes per kb, redundancy and completeness values, GC content, total length, number of contributing genomes, maximum number of paralogs, SCG clusters, and COG known our unknown functions, are represented. Gene clusters are identified as composing the core genome or as accessory and singletons.

Several DNA repair genes were detected among our three selected MAGs, mostly those involved with excision repair by the UVR system, mismatch repair and recombination (Figure 7). Within the UVR system, the UvrY and UvrD were detected only in DC_MAG_00005 (*Truepera sp*.). UvrB and UvrC were present in the three MAGs, and UvrA was not found only in DC_MAG_00021 (*Ca. Methylomirabilis*). Only DC_MAG_00005 (*Truepera sp*.) exhibited RecF, RecN and RecR were present in at least two MAGs, and RecO was found among the three MAGs. MutS was not detected only in DC_MAG_00016 (*Ca. Koribacter*). Other repair proteins were also predicted, such as LigD in DC_MAG_00021 (*Ca. Methylomirabilis*) and RadA in DC_MAG_00005 (*Truepera sp*.) and DC_MAG_00021 (*Ca. Methylomirabilis*). It was also detected among MAGs the predicted proteins *glucose starvation-inducible protein B* and *DNA protection during starvation protein* (Dps), both involved with starvation response. The predicted proteins involved with stress response were *general stress protein* (yocK), *peroxide-response repressor* (PerR), *superoxide dismutase* (SodA and SodB), and *universal stress protein* (TeaD).

**Figure 7.**
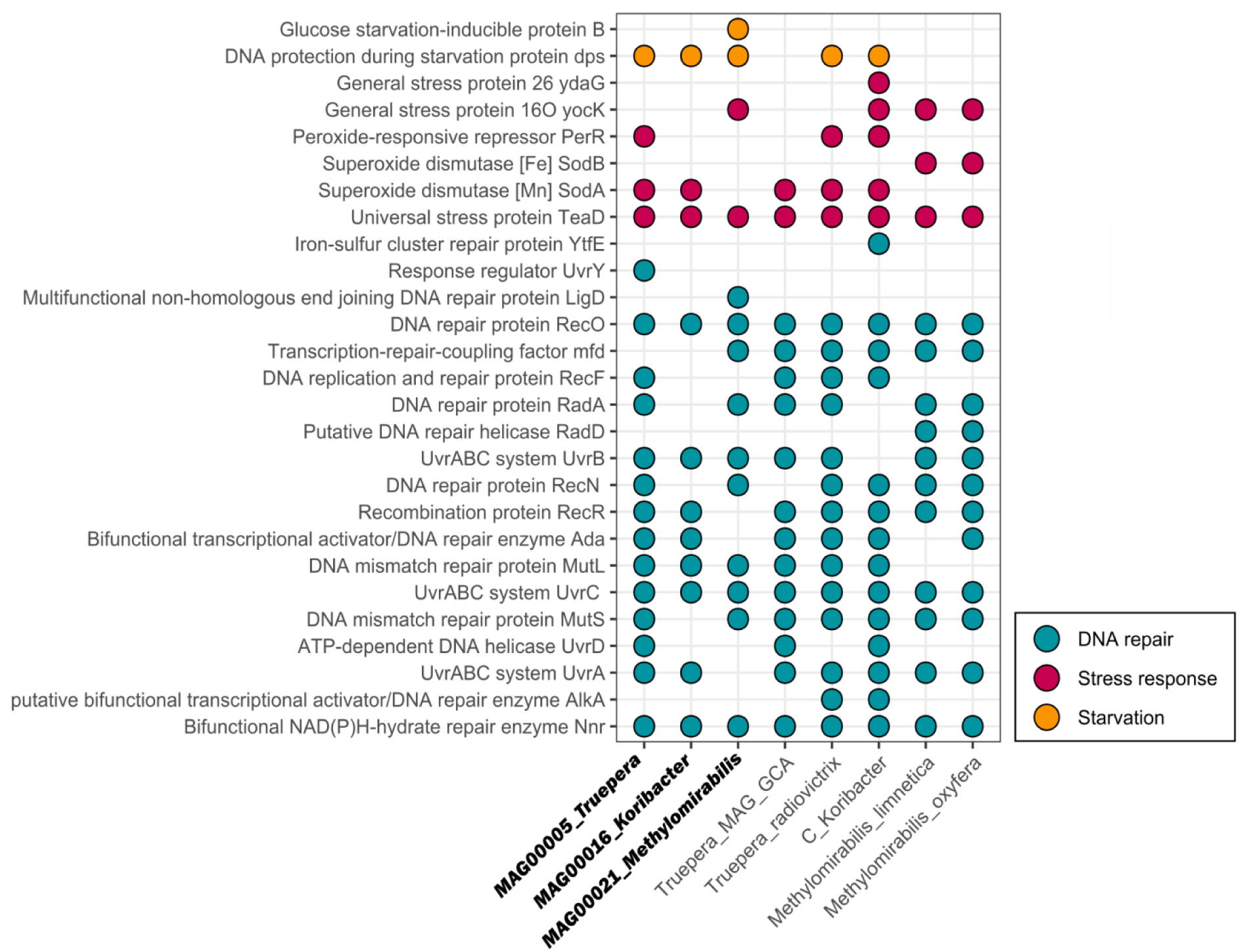
Representation of presence/absence of genes involved with DNA repair, stress and starvation responses, identified in the DC_MAG_00021 (*Ca. Methylomirabilis sp*.) DC_MAG_00005 (*Truepera sp*.) and DC_MAG_00016 (*Ca. Koribacter sp*.). The reference genomes from NCBI database used for pangenomic analysis are also represented, which include *Truepera radiovictrix* (DSM17093), *Truepera sp*. (GCA_002239005.1_ASM223900), *Ca. Koribacter versatilis* (NC_008009.1), *Ca. Methylomirabilis oxyfera* (FP565575.1) and *Ca. Methylomirabilis limnetica* (NZ_NVQC01000028.1).

## Discussion

The MAGs diversity identified in the present study allowed three approaches, addressed hereafter. First, the relationship between the *Monte Cristo* cave mineralogy, the physicochemical and the main taxonomic groups. The second approach concerns to the identification of MAGs and their metabolic potentials, mainly related to iron, sulfur, nitrogen and carbon elements, elucidating the functional diversity of *Monte Cristo* cave and adding new taxonomical data to the present knowledge about microbial assemblage in quartzite caves. Finally, the taxonomical, metabolic and ecological insights provided by the data set may serve as sources for astrobiology research, mainly concerning the habitability of extraterrestrial subsurface in rocky planets.

### Microbial genomes and the mineralogy of Monte Cristo cave

P1B was the site with the highest values of S (112.25 ppm), Mn (22.683 ppm), Cu (0.113 ppm) and conductivity (1424 µS/cm) in comparison to the other sites. The high conductivity in P1B may be related to the solubilization and remobilization of ions, in which several lineages related to bioleaching can be associated, especially by iron reduction and oxidation, as well as by sulfide oxidation. The MAGs recovered from P1B were associated with both heterotrophic and autotrophic metabolisms, including denitrification through nitrate reduction as one of the prevalent predicted processes. The majority of groups found in P1B (assigned within Actinobacteriota, Proteobacteria, Firmicutes, Bacteroidota, Patescibacteria and Deinococcota) were also detected in other karst caves, as Shuanghe Cave in China (Long et al., 2019) and Ozark cave in the USA (Oliveira et al., 2017), with exception of the Trueperaceae MAG (Deinococcota).

The P3 sample exhibited the lowest pH (4.4) detected within the cave, and intermediate values of Fe (176.43 ppm) and S (54.087 ppm). The only MAG recovered from P3 was assigned as *Mycobacterium sp*. A previous study has shown the abundance of *Mycobacterium* lineages in Moravian Karst (Czech Republic), which is likely explained by the presence of bat guano in the cave environment (Modra et al., 2017).

In the P7 site, it was found the highest values of Fe (227.993 ppm) and pH (7.15), and the lowest values of S (1.012 ppm) and conductivity (28 µS/cm). P7 was the only site under influence of a small stream in the surroundings. The prevalent lineages were classified within Nitrososphaeria, Chloroflexota, Acidobacteriota, Nitrospirota, Dormibacterota and Methylomirabilota. These groups were also described in other karst caves, such as in Chapada Diamantina (Brazil) (Marques et al., 2018), Ozark cave (USA) (Oliveira et al., 2017), Mizoram (India) (De Mandal et al., 2017) and Zhijin cave (China) (Dong et al., 2020). In this site, it was found the higher diversity of metabolic processes, mainly involved with chemolithotrophy, such as ammonia oxidation, sulfur oxidation, sulfate reduction, iron oxidation, nitrate reduction and anaerobic methane oxidation. This high metabolic versatility might reflect the presence of the small water flux near P7, which likely transport more nutrients and different geochemical compounds to this site, including from remobilization of minerals by microbial activity. Further, the detection of both aerobic and anaerobic microorganisms indicates a stratification condition probably due to the stream influence, which might create anoxic microniches a few centimeters below the surface. The presence of oxic and lowermost anoxic conditions within caves influenced by stratified sediments from a stream was reported in previous studies (Méndez-García et al., 2014; Reis et al., 2016).

Several groups of Archaea and Bacteria found in our samples have members related to bioleaching (the transformation of metals from ores into soluble metals). Different groups within Actinobacteriota, Proteobacteria, Nitrospirae and Firmicutes were previously related with iron mineral ores, especially acting in leaching of minerals such as iron oxides and sulfides in acid conditions. Some lineages from these phyla were reported acting in the iron oxidation (Hurtado et al., 2020). Under oxic conditions at neutral pHs, members of Actinobacteriota also can act in the reduction of iron (III) in iron-rich settings, including *Mycobacterium* (Zhang et al., 2019). In our study, *Mycobacterium* genomes were recovered in P1B and P3 under similar oxic conditions. Thus, supported by the high concentration of Fe in these points, our results suggest that this element may play a role in the energy metabolism of species within *Mycobacterium*. This genus, for example, is a known flocculating-flotating agent for hematite (Yang et al., 2007). Further, in the samples P1B and P7, genomes from the classes Gammaproteobacteria and Alphaproteobacteria were detected, which previously have had some of its members related with the processment of hematite (mineral found in P7). Also, according to a study made in subsurface by Reardon et al. (2004), Gammaproteobacteria and Alphaproteobacteria members were found attached to particles of a specular hematite medium, being the hematite-rich medium a possible attractive to the colonization of Fe(III)-reducers. At the P1B, rich in Fe and S, it was recovered a genome assigned within Xanthomonadales order, from Gammaproteobacteria class. This group was associated with acidic biofilms in pyrite mines in Germany (Ziegler et al., 2013), and although it is not directly involved with the leaching of the mineral, it was detected together with other groups that carry out the activity, probably being favored by a syntrophic relationship with these groups. Xanthomonadales was also detected in high abundance (∼25% of the total community) in acid mine drainage in the Appalachians (Grettenberger et al., 2017), simultaneously with iron-oxidizing members in conditions moderately acidic. They were also found associated with the suboxic and anoxic sediments along the Rio Tinto, Spain (Méndez-García et al., 2014), a highly metalliferous and acidic river considered an analog for the Martian subsurface.

Neubeck et al. (2017) presented the microbial diversity with 16S rRNA data from Chimaera ophiolite surface, a serpentinization-driven alkaline environment in Turkey. Among other groups, they found *Truepera* (Deinococcota), Rhyzobialles (Proteobacteria) and Xanthomonadales (Gammaproteobacteria) groups that were also found here at P1B. The P7 also have some common groups with the Chimaera ophiolite, such as Nitrospiraceae (Nitrospirae), Nitrososphaeria (Crenarchaeota) and Ktedonobacteraceae (Chloroflexota). Further, at Chimaera ophiolite the copper concentration was associated with the family Trueperaceae, that was also found in our P1B. At this point, Cu presented the highest concentration in comparison to the other sampling points.

The presence of some minerals, such as iron oxides and rutile, is another common information between Chimaera ophiolite and *Monte Cristo* cave, although closer comparisons are difficult due to the environmental contrasts between these ecosystems, such as the pH conditions and the geological setting. According to (Neubeck et al., 2017), the major diversity is outside the serpentinization zone, in which the alkaline conditions are higher. The authors found the Ktedonobacteraceae (Chloroflexota) in sites with low diversity and associated their occurrence with the nutrients limitatiher sampling points. They also associated the high indices of Fe with the adsorption of other elements, which contributed to the nutrient limitation and gave competitive advantage to microorganisms such as Ktedonobacteraceae. In fact, point P7 had the highest concentration of (Hoehler and Jørgensen, 2013; Müller and Hess, 2017) iron in comparison to the other points. Iron oxides form a complex with a large number of metallic cations, such as Pb^2+^, Cu^2+^, Ni^2+^, Co^2+^, Zn^2+^, and oxyanions such as Po_3_^4-^, SiO_4_^3-^, CrO_2_^4-^, AsO_3_^4-^ and MoO_2_^4-^. These complexes will create positive charges (adsorption of cations) and negative (adsorption of anions) on the mineral surface, which will increase the cationic and anionic exchange capacity of these minerals, influencing directly the chemical and physical behavior of the material (McKenzie, 1980; Tipping, 1981). Thus, the presence of abundant iron-rich crusts at our sampling sites may indicate the high adsorption of other elements, which contributes to the limitation of nutrients, and consequently, might explain the presence of different Ktedonobacteraceae genomes at the site, similarly to the observed for Chimaera ophiolite.

### Metabolic potential of MAGs and their relationships with the biogeochemical cycles

The survival of microorganisms depends mostly on its ability to obtain sufficient energy for cell maintenance and duplication (Hoehler and Jørgensen, 2013; Müller and Hess, 2017). In subsurface ecosystems, the microbial communities must rely on non-photosynthetic strategies in order to thrive under the energy-limiting condition. Here, it was possible to identified genes associated with a variety of metabolisms that allow these microbes to obtain carbon and energy from a wide range of sources. Three out of the six known CO2-fixation pathways were detected. The autotroph growth using Calvin-Benson-Bassham cycle (CBB) was found in one of our MAGs that was assigned to *Sulfuricella*, an anaerobic Betaproteobacteria that is known to oxidize sulfur and thiosulphate as its sole energy source (Kojima and Fukui, 2010; Watanabe et al., 2012). The Arnon–Buchanan reductive tricarboxylic acid (rTCA) cycle was also detected and it was associated with Nitrospira, which plays an important role in carbon and nitrogen cycles as an aerobic chemolithoautotrophic nitrite-oxidizing bacteria (Koch et al., 2019). Indeed it has been found in many microbial communities from caves, including the Kartchner Caverns, an extreme oligotrophic environment in Arizona, where the authors described microorganisms with diverse autotrophic potential and suggested that nitrite-oxidizing *Nitrospirae* represents noteworthy primary producers in this energy-limiting ecosystem (Ortiz et al., 2014; Tetu et al., 2013). Further, it was detected genes involved with the Wood–Ljungdahl pathway (WL) within *Ca. Bipolaricaulis*, a group of bacteria belonging to the yet-uncultivated phylum Bipolaricauliota, that has been recently described to harbor a high metabolic versatility (Hao et al., 2018; Youssef et al., 2019). The WL pathway may work in both reductive and oxidative directions, allowing the autotrophic (CO2-fixation) and the heterotrophic (fermentative metabolism) growth, which promotes an important advantage and versatility for microbial survival in harsh environmental conditions (Borrel et al., 2016; Müller et al., 2013).

Regarding the iron metabolism, only genes potentially involved with the oxidation of Fe (II) were identified. Iron is an essential element for almost all living organisms and, for some microbes, the oxidation of ferrous iron (Fe II) in ferric iron (Fe III) can power their metabolism(Bird et al., 2011). The *cyc2* was one of the most interesting gene related to iron oxidation found in our MAGs, since it has been previously described as a key component for iron oxidation in *Sphaerotilus lithotrophicus* (Emerson et al., 2013) and *Acidithiobacillus ferrooxidans* (Castelle et al., 2008), which are well-known neutrophilic and acidophilic Proteobacteria, respectively (Barco et al., 2015). In our study, *cyc2* was identified in an unclassified Proteobacteria and *Ca. Koribacter* (Actinobacteriota) MAGs. The use of iron as energy source by autotrophic Proteobacteria has also been reported in different cave ecosystems, such as in Mobile cave (Romania) (Kumaresan et al., 2018) and in Tjuv Ante’s cave (Sweden) (Mendoza et al., 2016). Our results suggest that iron-oxidizing bacteria might be involved with primary production of the *Monte Cristo* cave ecosystem as well.

In addition, *Monte Cristo* cave showed to harbor a variety of microorganisms related to both oxidative and reductive pathways within sulfur metabolism, including sulfur and thiosulfate oxidation (*sox* genes), sulfate reduction (*sat*) and sulfide oxidation (*sqr*). Several studies have reported that sulfur-oxidizing bacteria play an important role as primary producers in different cave ecosystems, together with ammonia- and nitrite-oxidizing microorganisms (e.g. Chen et al., 2009; Macalady et al., 2008; Marques et al., 2018; Zhao et al., 2017). The sulfur-oxidizing genes identified in our MAGs were related to Nitrospiraceae and Betaproteobacteriales. This sulfur oxidation occurs in oxic conditions and likely leads to the production of sulfate, which can be then reduced to sulfite and sulfide in anoxic zones by the sulfate reducers (Zerkle et al., 2016). In fact, it was possible to reconstruct genomes of microorganisms potentially involved with this process (by the identification of *sat* genes), such as unclassified Gammaproteobacteria and Nitrospiraceae. Some of our MAGs also possess the *sqr* gene, which can potentially oxidize the sulfide produced by the sulfate reducers. These oxidative and reductive sulfur pathways might be occurring in oxic and anoxic microniches created by the stratified sediments attached to the cave wall, and suggests a syntrophic relationship occurring among these microorganisms. Further, genomes assigned as anaerobic methane oxidizers were also detected in *Monte Cristo* cave and they could potentially maintain a syntrophic relation with the sulfate-reducing bacteria within anoxic strata, as was observed for anchialine caves by Pohlman (2011).

Additionally, nitrogen is often shown as a key element for the energy metabolism within caves ecosystems (Ortiz et al., 2014; Wiseschart et al., 2019). Although it was not possible to detect genes for nitrogen fixation (nitrogenase - *nifH*), a genome assigned to the known nitrogen-fixing *Bradyrhizobium sp*. (DC_MAG_00058) was assembled. Interestingly, the results also showed evidence for the complete pathway of microbial-mediated denitrification by identifying the genes required for the reduction of nitrate (*nar*), nitrite (*nir*), nitric oxide (*nor*) and nitrous oxide (*nos*). It was also noticed that these genes were detected among diverse groups including cultivated and yet-uncultivated bacteria such as Actinobacteria (*Mycobacterium* and *Micolicibacter*), *Ca. Bipolaricaulis* and *Flaviosilibacter*. Further, two MAGs were assigned to well characterized ammonia-oxidizing archaea (AOA) within Crenarchaeota (*Ca. Nitrosotalea*) and Thaumarchaeota (*Nitrososphaera*). The AOA have already been proposed as important members to the energy dynamics of oligotrophic karstic environments, playing a role in aerobic nitrification (Ortiz et al., 2014; Tetu et al., 2013). Indeed, it is suggested that Archaea, especially Thaumarchaeota, are well adapted to low-nutrient environments, and our findings contribute to corroborate it (Bates et al., 2011; Martens-Habbena et al., 2009).

The diversity of chemolithoautotrophic and chemoorganotrophic pathways described in our genomes suggests the maintenance of complex ecological processes within terrestrial aphotic ecosystems as *Monte Cristo* cave. However, it is important to highlight that the direct relationships between microorganisms and the biogeochemical cycles occurring in *Monte Cristo* cave would require further studies and other techniques to measure microbial activity, such as metatranscriptomic and stable isotope approaches.

### Comparison and survival strategies of Truepera sp., Ca. Methylomirabilis sp. and Ca. Koribacter sp. MAGs

Although the majority of groups classified from our genomes has been described in different karst caves worldwide, the Trueperaceae (DC_MAG_00005) was the only which, to the best of our knowledge, has not been previously detected in these ecosystems. This chemoorganotroph family is known for its ionizing radiation resistance and was mostly obtained from geothermal (Albuquerque et al., 2005), saline lake (Lavrentyeva et al., 2020) and dry environments as soil crusts (Weber et al., 2018). The first isolates showed an optimal growth at 50°C, a growth up to 6% of NaCl and a capability to survive (60% of the cells) to 5.0 kGy (Albuquerque et al., 2005). Our Trueperaceae MAG (DC_MAG_00005) exhibited several genes involved with different DNA repair mechanisms, oxidative stress response and starvation. The majority of these genes is also found in the reference genomes, with the exception of UvrY. Although the cave system has no radiation exposure, other conditions such as high salinity or desiccation might be capable of selecting radiation resistance microorganisms, since the cell repair processes for these parameters are often the same (Musilova et al., 2015). Indeed, the Trueperaceae MAG (DC_MAG_00005) was recovered from the P1B site, which corresponds to a dry rock sampled from the cave wall and presented the highest values of conductivity, indicating a higher salt concentration in comparison with the other sites. Our pangenomic analysis suggested that our Trueperaceae MAG (DC_MAG_00005) might represent a novel genus or even a novel family within the order Deinococcales.

Another interesting genome recovered from *Monte Cristo* cave was DC_MAG_00021 assigned as *Ca. Methylomirabilis*. This microorganism was firstly discovered in Nullarbor Cave (Australia) as NC-10 phylum (Holmes et al., 2001), and further genomic and isotopic studies have described them as the only known organism that can couple anaerobic methane oxidation to the reduction of nitrite to N2 (Ettwig et al., 2010; Versantvoort et al., 2018). Both anaerobic and aerobic methanotrophic metabolism was observed in previous studies from submerged portions of Nullarbor Cave (Australia) (Holmes et al., 2001), Sulzbrunn Cavern (Germany) (Karwautz et al., 2018), Movile Cave (Romania) (Kumaresan et al., 2018) and Heshang Cave (China) (Zhao et al., 2018). Our samples are not associated with submerged aquatic systems and the presence of anaerobic methane-oxidizing microorganisms on the surface of the cave wall might be explained by a potential stratification with anaerobic microniches lowermost the oxic surface strata, as observed in previous studies (Méndez-García et al., 2014; Reis et al., 2016). The DC_MAG_00021 (*Ca. Methylomirabilis*) exhibited several DNA repair genes, and was the only genome which presented the coding-gene for LigD, a protein ligase involved with non-homologous end joining (NHEJ) DNA repair caused by DNA double-strand breaks (Shuman and Glickman, 2007). General stress response and starvation proteins were also found in this MAG, indicating a molecular capability of surviving under stressful conditions, likely oligotrophy. These starvation proteins were not detected in the reference genomes of *Ca. Methylomirabilis*. The comparison between our *Ca. Methylomirabilis* with the reference genomes indicates our MAG as likely a novel taxon within Methylomirabilaceae and further phylogenomic analysis is needed for its description.

The *Ca. Koribacter* (DC_MAG_00016) was the third genome selected for pangenomic analysis due to the presence of genes related to iron oxidation. A recent study suggests a Mn oxidation capacity for this genus due to the identification of four proteins homologous to multicopper oxidases, which are known to perform this metabolism (Allward et al., 2018). However, these proteins were not detected in our MAG. Otherwise, the *cyc2* e sulfocyanin genes were the two functions related to iron oxidation found in this MAG, which suggests a potential role in iron oxidation. Interestingly, the copper protein sulfocyanin was found mostly in Archaea, and combined with cytochrome *cbb*3 oxidase, seem to be involved in the electron transfer chain between Fe(II) and O_2_ (Ilbert and Bonnefoy, 2013). However, despite the presence of these genes, the iron oxidation capacity of *Ca. Koribacter* remains unclear. Concerning the DNA repair genes in our DC_MAG_00016, *uvrB* (that encodes nucleotide excision repair protein) was the only that was not detected in the reference genome. In this same MAG, the nitric oxide reductase (*norC*) and *sat* genes were identified, which are involved with denitrification and sulfate reduction, respectively. While nitric oxide reduction was previously described in *Ca. Koribacter versatilis* (GCA_000014005.1), the *sat* was not detected in this reference genome. According to (Ward et al., 2009), *Ca. Koribacter* is also capable of CO oxidation (through *cox* genes that were also found in our *Ca. Koribacter* MAGs) and degradation of complex polymers, which is speculated that this capacity arose as scavenging strategies to optimize life in low-carbon environments. Indeed, *Koribacter* sequences were found in iron-rich and oligotrophic environments (Dedysh and Damsté, 2018; Kielak et al., 2016; Ward et al., 2009). Nevertheless, their physiological properties and ecological roles in these ecosystems remains unclear.

### Astrobiological implications

Currently on Mars, potential lifeforms would be challenged by environmental conditions as low atmospheric pressure, extreme temperature variations, strong UV and cosmic radiation, high concentrations of heavy metals, high pH, a CO_2_-dominated atmosphere and low concentrations of organic compounds (Nixon et al., 2013; Westall et al., 2013). The lifeforms best adapted to those conditions would be related to chemolithotrophic metabolisms, likely using inorganic compounds for reductive and oxidative reactions, such as methane, H_2_, iron and sulfate, and CO_2_ as a carbon source (Nixon et al., 2013; Seto et al., 2019). Some aspects identified in the present study may be of interest for astrobiological prouposes and may shed light in these presented questions. In terms of interaction between microorganisms and minerals, the occurrence of microbiota in samples containing hematite are especially interesting due the possibility that this mineral plays a central role as electron donor or acceptor for metabolic processes (Nixon et al., 2013). The recognition of metabolic pathways related to methane is also of interest due their application to understand atmospheric variation of this gas during the Martian year (Webster et al., 2018). Also concerning metabolic pathways, the three selected genomes with metabolic and survival distinguish features (*Truepera sp*., *Ca. Methylomirabilis sp*. and *Ca. Koribacter sp*.), provide models of molecular adaptation of present life in the extreme environments outside Earth. Lastly, the identification of Martian caves opens new perspectives for modern life refugies in response to the harsh surface conditions of Mars’ surface (Cushing and Titus, 2010), and there is still need of identifying potential analogue environments on Earth for such Martian systems.

*In situ* (e.g. Baird et al., 1976; Bish et al., 2013; Johnson et al., 2016; Rieder et al., 1997; Squyres et al., 2009; Vaniman et al., 2014) and orbital analysis (e.g. Christensen et al., 2003; Singer et al., 1979) widely indicated the common occurrence of ferric oxides and oxyhydroxides, mainly magnetite and ilmenite as primary minerals (from past igneous activity), and hematite, goethite, akaganeite and ferrous carbonate as secondary minerals (from weathering and alteration processes), being the last ones the components of the reddish, fine-grained (<5 μm) dust over the surface of Mars (Ehlmann and Edwards, 2014 and references therein). The secondary iron mineral phases are related to the groundwater processes and diagenesis (Squyres et al., 2009), occurred from ca. 3.5 billion years ago, during the Hesperian period, onwards (Ehlmann and Edwards, 2014). This iron abundance over Mars’ surface and bedrocks would be useful for metabolism, being the Fe^2+^ acting as electron donors, and Fe^3+^as electron acceptors. However, both metabolic pathways depend on the recognition of other compounds, such as oxygen or nitrate for iron and sulfur oxidation, and organic molecules from extraterrestrial input, and H_2_ for the reducing iron and sulfur metabolisms. Also, CO would be useful for Fe- and S-reduction metabolism, and this gas has been already detected in the Martian atmosphere, leading to question if its abundance may be an indicative of absence of life (Nixon et al., 2013). In addition, the presence of silicates on Mars crust, as olivine and pyroxene, can favor an iron oxidation metabolism, which on Earth is performed by several microbial lineages that use O_2_ or nitrate as electron acceptors, and potentially by novel lineages that are being recently discovered in different ecosystems, such as the *Ca. Koribacter sp*. detected in our samples (Dedysh and Damsté, 2018). This data demonstrates that the study of poorly explored environments on Earth, such as the *Monte Cristo* cave, is crucial to reveal novel ecological roles and capacities of these lineages. In Addition to the recognition of novel pathways, the recognition of ecological constraints and fostering of metabolisms like this of *Ca. Koribacter* may be interesting for elucidating possible extant metabolisms on Mars, such as oxidation of iron compounds using CO2 as carbon sources (Nixon et al., 2013).

Since the presence of methane was detected by telescopic, orbital and landed missions since the 1990s (Webster et al., 2018), questions concerning its biological origin have been raised. On Earth, this molecule presents an intimate relationship with life, mainly through methanogenic processes due its role as electron donor. Atreya et al. (2007) and Hu et al. (2016) discussed the possible processes involved in this seasonal methane increase in the Martian atmosphere during the end of summer in the northern hemisphere (Webster et al., 2018) and pointed that microorganisms may be the responsible for this phenomena by converting organic matter into methane in presence of a liquid solution. On the other hand, examples of sinks that promote the methane to return to autumn-winter levels may be attributed to UV photodissociation (Atreya et al., 2007), strong reactions to surface oxidants (Atreya et al., 2007), horizontal advection and diffusion (Hu et al., 2016), sequestration by recently discovered methyl compounds exposed to surface by wind erosion (Jensen et al., 2014) and electron drift motion in electric fields promoted by dust storms (Farrell et al., 2006). No biological consumption has been proposed. However, it does not mean that life is not involved in the Martian methane cycle. On the contrary: sazonal abundance of methane may act as an episodic niche to be explored, however, not reaching levels enough to cause the global atmospheric methane concentration to return to autumn-winter concentrations. Also, punctual methane sources, such as areas subject to serpentinization, may play an interesting role as micro and restricted niches for methane-based ecosystems. In this sense, methane consumption metabolism is a central piece, and oxidation pathways under a reductive atmosphere may offer a plausible explanation. Anaerobic methane oxidation performed by bacteria as *Ca. Methylomirabilis sp*., explored in this research, is a promising model that assembles two important aspects of the methane tale: it consumes methane in anaerobic conditions and occurs in subsurface (Versantvoort et al., 2018). Thus, lineages of the group including *Ca. Methylomirabilis* shows a potential metabolism to be explored on Mars (Seto et al., 2019).

Although subsurface ecosystems on Mars are shielded from UV and cosmic radiation, the need for a radiation resistance capacity of potential lifeforms is critical when one considers about their persistence for long periods, since they might be eventually exposed to aeolian dispersion processes (as occurs on Earth) that might require a strong molecular adaptation for their survival under the high levels of radiation on Mars’ surface (Azua-Bustos et al., 2019). Further, the radiation resistance mechanisms in microbes on Earth are also associated with the survivability under desiccation and high salinity conditions, which are two strong selective pressures for potential lifeforms in Mars’ subsurface. Indeed, at *Monte Cristo* cave was detected a bacteria classified as *Truepera sp*., which is known for its strong radiation resistance (Albuquerque et al., 2005), showing that subsurface ecosystems on Earth might also select this type of adaptation.

Questions concerning the limits and constraints imposed to a possible extant life in the Martian surface have been repeatedly addressed, mainly due the extreme conditions it imposes to the cell integrity, including possible physical damage (e.g. meteoroid bombardment, dust storms, extreme temperature variations) and chemical injuries (e.g. solar flares, UV radiation, and high-energy particles input from space) (de Vera et al., 2019; Mazur et al., 1978; Westall et al., 2013). As pointed by Mazur et al. (1978), the idea of oasis offers an environmental alternative for an Earthly-like life to exist, and, in this sense, caves may be one of these environments (Boston et al., 2001; Cushing and Titus, 2010), where life is supposed to be protected from extreme environmental variations imposed to the Martian surface. The case study of *Monte Cristo* cave offers an important window for the oasis hypothesis for life on Mars: a wide range of metabolism pathways, exploring different elements in absence of light, potentially involved in syntrophic relationships, and protected from variable conditions of surface. Also, the study of such restricted environments offers opportunities to elaborate minimum-impact protocols of exploration and optimization of operational aspects, as advocated by Boston et al. (2001).

Finally, the diversity of metabolic potentials and survival strategies of microbes from the *Monte Cristo* cave offers a window to observe and understand life adapted to oligotrophic conditions of limited light offering to totally dark. This condition, associated with an environment rich in iron and silica, as expected for the environments from other bodies in the Solar System, make ecosystems like this from *Monte Cristo* cave important for astrobiology and its interest in habitability of modern life.

## Conclusion

The oligotrophic conditions identified in the *Monte Cristo* cave, mainly related to the limitation for accessing light, associated to the presence of silica, iron, manganese and sulfur, make this environmental setting an important opportunity to understand life under limitant and stressful conditions, suitable for the discussion concerning the habitability of modern life in other Solar System surfaces and subsurfaces, such as Mars. The quartzite *Monte Cristo* cave provided important taxonomical, life strategy and ecological information for the search for extant life in future missions on Mars and other silica and iron-rich surfaces. All this contribution is based on the genomic reconstruction and metagenomic assembly here presented, and their relationship to the physicochemical and cave mineral composition, which comprises seven different mineral phases. The relationship between the microbiota and the biotope concerns i) mainly the abundance of iron and water are direct related to a greater number of metabolic pathways (P7); and ii) putative occurrence of bioleaching by the microorganisms, which, in turn, use the remobilized metal in their metabolism. The metabolic pathways identified in the *Monte Cristo* cave comprises mainly aerobic and anaerobic chemolithotrophic activity related to oxidation of iron, methane, sulfur, ammonia, as well the reduction of sulfate several nitrogen compounds, showing versatil microbial metabolisms, mainly in face of adaptive demand under oligotrophic conditions. Microbial taxa comprises Bacteria and Archaea classified within 13 phylum including Actinobacteriota, Proteobacteria, Acidobacteriota, Crenarchaeota, Bacteroidota, Firmicutes, Nitrospirota, Chloroflexota, Patescibacteria, Bipolaricaulota, Dadabacteria, Deinococcota, Methylomirabilota, as well as unclassified bacterial phylum, which include many lineages able to bioleaching, especially metals such as iron oxides and sulfides. Finally, the analysis of the quartzite *Monte Cristo* cave shed light in different aspects of a poor-known microbiota of these ecosystems, improving our knowledge about this type of life diversity and opening perspectives for further studies.

## Author Contributions

All the authors interpreted the results, wrote and reviewed the manuscript, and approved the submitted version. AB, FC, ES, AV, FR, DG collected the samples, conceived and designed the experiments and analyzed the data. AB and MA performed bioinformatic analysis, analyzed the data and prepared figures and tables. DG coordinated the study and provided the financial support for the experiments. VP contributed to reagents/materials/analysis tools. FC, ES, AV, GB and VT performed geological description and mineralogical analysis.

## Acknowledgements

We thank the Deep Carbon Observatory’s Census of Deep Life initiative for the support of metagenome sequencing. We are very grateful for the support of Frederick Colwell and the assistance of Mitch Sogin, Joseph Vineis, Andrew Voorhis, and Hilary Morrison at MBL (Woods Hole, MA, USA).

## Supplementary material

Supplementary Figure 1. General metabolic potential of the 61 medium- and high-quality genomes are represented as absolute abundance. Genes related to metabolisms of carbon, nitrogen, and sulfur, as well as hydrogenases, were selected by MetabolisHMM tool. Genes involved with the iron cycle were selected by the FeGenie tool. The taxonomy is represented until the deepest classified level according to GTDB-Tk.

Supplementary Table 1. Information about the 61 medium- and high-quality genomes, including the taxonomy classification by KEGG and the GTDB-Tk, the total length, number of contigs, N50, GC content, and the values of completeness and redundancy.

